# Dog Size and Patterns of Disease History Across the Canine Age Spectrum: Results from the Dog Aging Project

**DOI:** 10.1101/2022.05.03.490110

**Authors:** Yunbi Nam, Michelle White, Elinor K. Karlsson, Kate E. Creevy, Daniel Promislow, Robyn L. McClelland, The Dog Aging Project Consortium

## Abstract

Age in dogs is associated with the risk of many diseases, and canine size is a major factor in that risk. However, the size effect is not as simple as the age effect. While small size dogs tend to live longer, some diseases are more prevalent among small dogs. Utilizing owner-reported data on disease history from a substantial number of companion dogs, we investigate how body size, as measured by weight, associates with the prevalence of a reported condition and its pattern across age for various disease categories. We found significant positive associations between weight and prevalence of skin, bone/orthopedic, gastrointestinal, ear/nose/throat, cancer/tumor, brain/neurologic, endocrine, and infectious diseases. Similarly, weight was negatively associated with the prevalence of eye, cardiac, liver/pancreas, and respiratory disease categories. Kidney/urinary disease prevalence did not vary by weight. We also found that the association between age and disease prevalence varied by dog size for many conditions including eye, cardiac, orthopedic, ear/nose/throat, and cancer. Controlling for sex, purebred/mixed breed, and geographic region made little difference in all disease categories we studied. Our results align with the reduced lifespan in larger dogs for most of the disease categories but suggest potential avenues for further examination.

## Introduction

Age is the single greatest predictor of disease risk for most causes of mortality in both human and dog populations (1–3). However, unlike in humans, for many dog diseases, body size is a comparably important predictor of risk (1,4,5). Companion dogs show considerable variation in longevity across size classes (6–8). Between species of mammals, larger ones tend to live longer than smaller ones, while within species, smaller individuals tend to live longer than larger individuals (9). Accordingly, dogs from larger size classes tend to have a shorter lifespan. Additionally, different size classes of dogs tend to manifest with different diseases, and ultimately to die from different causes. For instance, larger breed dogs more often die of musculoskeletal and gastrointestinal causes whereas smaller dogs die more frequently of endocrine causes (1,2).

Jin et al (10) studied a multimorbidity index (e.g. a count of the number of conditions a dog has experienced) and found that although the index increased steadily with age, the size of the dog had no significant effect on the index. This suggests that over time, dogs in larger size classes are not accumulating *more* conditions, but rather *different* conditions. An understanding of which conditions manifest differently across age and size could inform our understanding of size-related longevity differences. The Dog Aging Project (DAP) provides a unique opportunity to address these issues in a large community-based population of companion dogs. Specifically, we use data from the “DAP Pack”, a collection of over 25,000 survey respondents from across the US.

## Methods

The Dog Aging Project (DAP) was launched in Fall 2019 (11). Dog owners self-selected and could enroll one dog per household. Enrollment consisted of completing a web-based Health and Life Experience Survey (HLES) that elicited from the dog owner information on a wide array of topics. The curated 2020 release of the HLES data contained 27,541 survey records collected on or before December 31st, 2020. The survey consisted of ten sections including dog demographics, environment, health status, and owner demographics. In the health status section, dog owners were asked if their dogs have acquired and been diagnosed with various medical conditions, organized by organ system or pathophysiologic process category. We focused on thirteen disease categories of interest within which conditions were reported in 500 or more dogs. Categories that were considered include skin disorders, infectious or parasitic disease, orthopedic, GI, ocular, ear/nose/throat, kidney/urinary, cancer, cardiac, neurologic, liver/pancreas, respiratory, and endocrine disorders.

We presented the frequency and proportion of owner-reported disease history by age categories in Table 1, and by weight categories in Table 2. Disease categories are listed in descending order of overall frequency. Owners were asked to answer the year and month the dog was born if known or to provide an estimated age. Based on this calculated age, we categorized them into puppies (<1 year), adolescents (1 to <3 years), young adults (3 to <7 years), older adults (7 to <11 years), or seniors (11+ years). Owners were asked to report the dog’s exact weight in pounds, to the best of their knowledge. We converted the unit of weight from pounds to kilograms and classified them into five size groups: <10 kg, 10 to <20 kg, 20 to <30 kg, 30 to <40kg, or 40+ kg.

**Table 1:** Proportion with Disease History by Age

|  | Puppy<br>(<1yr) | Adolescent<br>(1 to <3yr) | Young Adult<br>(3 to <7yr) | Older Adult<br>(7 to <11yr) | Senior<br>(≥11yr) | Overall |
| --- | --- | --- | --- | --- | --- | --- |
| Disease | N=591 | N=4622 | N=8249 | N=7666 | N=6413 | N=27541 |
| Skin | 40 (7%) | 796 (17%) | 2181 (26%) | 2556 (33%) | 2342 (37%) | 7915 (29%) |
| Infection/Parasites | 147 (25%) | 1269 (27%) | 2291 (28%) | 2009 (26%) | 1623 (25%) | 7339 (27%) |
| Bone/Orthopedic | 7 (1%) | 239 (5%) | 849 (10%) | 1699 (22%) | 2493 (39%) | 5287 (19%) |
| Gastrointestinal | 39 (7%) | 488 (11%) | 1059 (13%) | 1158 (15%) | 1170 (18%) | 3914 (14%) |
| Eye | 31 (5%) | 283 (6%) | 586 (7%) | 906 (12%) | 1819 (28%) | 3625 (13%) |
| Ear/Nose/Throat | 19 (3%) | 302 (7%) | 745 (9%) | 916 (12%) | 1587 (25%) | 3569 (13%) |
| Kidney/Urinary | 14 (2%) | 153 (3%) | 406 (5%) | 591 (8%) | 957 (15%) | 2121 (8%) |
| Cancer/Tumors | 1 (<1%) | 21 (<1%) | 177 (2%) | 586 (8%) | 966 (15%) | 1751 (6%) |
| Cardiac | 3 (<1%) | 45 (<1%) | 167 (2%) | 440 (6%) | 912 (14%) | 1567 (6%) |
| Brain/Neurologic | 2 (<1%) | 25 (<1%) | 188 (2%) | 359 (5%) | 750 (12%) | 1324 (5%) |
| Liver/Pancreas | 2 (<1%) | 24 (<1%) | 125 (2%) | 277 (4%) | 542 (8%) | 970 (4%) |
| Respiratory | 4 (<1%) | 53 (1%) | 132 (2%) | 242 (3%) | 519 (8%) | 950 (3%) |
| Endocrine | 0 (<1%) | 8 (<1%) | 77 (<1%) | 310 (4%) | 518 (8%) | 913 (3%) |

**Table 2:** Proportion with Disease History by Weight Category

|  | <10kg | 10 to <20kg | 20 to <30kg | 30 to <40kg | ≥40kg | Overall |
| --- | --- | --- | --- | --- | --- | --- |
| Disease | N=6207 | N=5613 | N=8219 | N=5167 | N=2335 | N=27541 |
| Skin | 1628 (26%) | 1531 (27%) | 2369 (29%) | 1621 (31%) | 766 (33%) | 7915 (29%) |
| Infection/Parasites | 1186 (19%) | 1593 (28%) | 2441 (30%) | 1478 (29%) | 641 (27%) | 7339 (27%) |
| Bone/Orthopedic | 1187 (19%) | 931 (17%) | 1461 (18%) | 1181 (23%) | 527 (23%) | 5287 (19%) |
| Gastrointestinal | 892 (14%) | 786 (14%) | 1119 (14%) | 749 (14%) | 368 (16%) | 3914 (14%) |
| Eye | 1079 (17%) | 800 (14%) | 934 (11%) | 571 (11%) | 241 (10%) | 3625 (13%) |
| Ear/Nose/Throat | 827 (13%) | 713 (13%) | 915 (11%) | 752 (15%) | 362 (16%) | 3569 (13%) |
| Kidney/Urinary | 530 (9%) | 452 (8%) | 639 (8%) | 369 (7%) | 131 (6%) | 2121 (8%) |
| Cancer/Tumors | 273 (4%) | 327 (6%) | 565 (7%) | 428 (8%) | 158 (7%) | 1751 (6%) |
| Cardiac | 671 (11%) | 362 (6%) | 301 (4%) | 177 (3%) | 56 (2%) | 1567 (6%) |
| Brain/Neurologic | 362 (6%) | 278 (5%) | 346 (4%) | 235 (5%) | 103 (4%) | 1324 (5%) |
| Liver/Pancreas | 393 (6%) | 209 (4%) | 195 (2%) | 130 (3%) | 43 (2%) | 970 (4%) |
| Respiratory | 385 (6%) | 169 (3%) | 207 (3%) | 124 (2%) | 65 (3%) | 950 (3%) |
| Endocrine | 194 (3%) | 190 (3%) | 264 (3%) | 176 (3%) | 89 (4%) | 913 (3%) |

**Table 3:** Association of Age and Weight with Lifetime Prevalence (Part 1: Conditions More Common in Larger Dogs)

| Condition | Characteristic | Model 1<sup>a</sup> |  |  | Model 2<sup>b</sup> |  |  | Model 3<sup>c</sup> |  |  |
| --- | --- | --- | --- | --- | --- | --- | --- | --- | --- | --- |
|  |  | PR | 95% CI | p-value | PR | 95% CI | p-value | PR | 95% CI | p-value |
| Skin | Age | 1.29 | (1.27, 1.31) | <0.001 | 1.29 | (1.27, 1.31) | <0.001 | 1.27 | (1.25, 1.3) | <0.001 |
|  | Weight | 1.12 | (1.1, 1.14) | <0.001 | 1.12 | (1.1, 1.14) | <0.001 | 1.13 | (1.11, 1.15) | <0.001 |
|  | Age * Weight | - | - | - | 0.98 | (0.96, 1) | 0.02 | 0.98 | (0.96, 0.99) | <0.01 |
| Bone/Orthopedic | Age | 2 | (1.95, 2.04) | <0.001 | 2 | (1.96, 2.05) | <0.001 | 1.98 | (1.93, 2.03) | <0.001 |
|  | Weight | 1.22 | (1.19, 1.25) | <0.001 | 1.17 | (1.13, 1.2) | <0.001 | 1.17 | (1.13, 1.2) | <0.001 |
|  | Age * Weight | - | - | - | 1.08 | (1.05, 1.11) | <0.001 | 1.08 | (1.05, 1.11) | <0.001 |
| Gastrointestinal | Age | 1.22 | (1.19, 1.26) | <0.001 | 1.22 | (1.19, 1.26) | <0.001 | 1.21 | (1.18, 1.25) | <0.001 |
|  | Weight | 1.04 | (1.01, 1.07) | 0.01 | 1.04 | (1.01, 1.07) | 0.01 | 1.03 | (1, 1.06) | 0.06 |
|  | Age * Weight | - | - | - | 0.98 | (0.96, 1.01) | 0.24 | 0.99 | (0.96, 1.02) | 0.42 |
| Ear/Nose/Throat | Age | 1.76 | (1.7, 1.82) | <0.001 | 1.74 | (1.68, 1.79) | <0.001 | 1.73 | (1.68, 1.79) | <0.001 |
|  | Weight | 1.16 | (1.12, 1.19) | <0.001 | 1.21 | (1.17, 1.24) | <0.001 | 1.19 | (1.16, 1.23) | <0.001 |
|  | Age * Weight | - | - | - | 0.89 | (0.87, 0.92) | <0.001 | 0.9 | (0.87, 0.92) | <0.001 |
| Cancer/Tumors | Age | 2.56 | (2.45, 2.67) | <0.001 | 2.54 | (2.43, 2.66) | <0.001 | 2.52 | (2.41, 2.64) | <0.001 |
|  | Weight | 1.4 | (1.35, 1.45) | <0.001 | 1.33 | (1.27, 1.39) | <0.001 | 1.35 | (1.28, 1.41) | <0.001 |
|  | Age * Weight | - | - | - | 1.07 | (1.03, 1.11) | <0.001 | 1.07 | (1.02, 1.11) | <0.01 |
| Brain/Neurologic | Age | 2.46 | (2.33, 2.59) | <0.001 | 2.48 | (2.35, 2.62) | <0.001 | 2.49 | (2.35, 2.63) | <0.001 |
|  | Weight | 1.08 | (1.02, 1.15) | <0.01 | 1.04 | (0.97, 1.12) | 0.3 | 1.01 | (0.94, 1.09) | 0.7 |
|  | Age * Weight | - | - | - | 1.06 | (1, 1.12) | 0.05 | 1.07 | (1.02, 1.13) | 0.01 |
| Endocrine | Age | 2.6 | (2.46, 2.75) | <0.001 | 2.6 | (2.46, 2.75) | <0.001 | 2.59 | (2.45, 2.74) | <0.001 |
|  | Weight | 1.27 | (1.19, 1.35) | <0.001 | 1.26 | (1.18, 1.35) | <0.001 | 1.24 | (1.16, 1.33) | <0.001 |
|  | Age * Weight | - | - | - | 1.01 | (0.96, 1.06) | 0.74 | 1.02 | (0.97, 1.07) | 0.52 |
| Infection/Parasites | Age | 0.97 | (0.95, 0.99) | <0.01 | 0.97 | (0.96, 0.99) | 0.01 | 0.96 | (0.94, 0.98) | <0.001 |
|  | Weight | 1.09 | (1.07, 1.11) | <0.001 | 1.1 | (1.08, 1.12) | <0.001 | 1.11 | (1.09, 1.13) | <0.001 |
|  | Age * Weight | - | - | - | 1.04 | (1.02, 1.06) | <0.001 | 1.04 | (1.02, 1.06) | <0.001 |

*Note:*
Age (years) and Weight (kg) are standardized by subtracting their means 7, 23 and dividing by their stadard deviations 4, 13.
<sup>a</sup> A model with the main effects of age and weight.
<sup>b</sup> A model with the main effects of age, weight, and the interaction.
<sup>c</sup> A model with the main effects of variables in Model 2 plus adjusted for sex, purebred/mixed breed, and geographic region.

**Table 4:** Association of Age and Weight with Lifetime Prevalence (Part 2: Conditions Less or Equally Common in Larger Dogs)

| Condition | Characteristic | Model 1<sup>a</sup> |  |  | Model 2<sup>b</sup> |  |  | Model 3<sup>c</sup> |  |  |
| --- | --- | --- | --- | --- | --- | --- | --- | --- | --- | --- |
|  |  | PR | 95% CI | p-value | PR | 95% CI | p-value | PR | 95% CI | p-value |
| Eye | Age | 1.86 | (1.81, 1.92) | <0.001 | 1.8 | (1.74, 1.86) | <0.001 | 1.81 | (1.75, 1.87) | <0.001 |
|  | Weight | 0.93 | (0.9, 0.96) | <0.001 | 0.98 | (0.94, 1.02) | 0.27 | 0.96 | (0.93, 1) | 0.05 |
|  | Age * Weight | - | - | - | 0.89 | (0.87, 0.92) | <0.001 | 0.91 | (0.88, 0.94) | <0.001 |
| Kidney/Urinary | Age | 1.76 | (1.69, 1.83) | <0.001 | 1.75 | (1.68, 1.82) | <0.001 | 1.72 | (1.65, 1.79) | <0.001 |
|  | Weight | 1.01 | (0.97, 1.05) | 0.77 | 1.02 | (0.98, 1.07) | 0.32 | 1.07 | (1.02, 1.12) | <0.01 |
|  | Age * Weight | - | - | - | 0.96 | (0.93, 1) | 0.06 | 0.94 | (0.91, 0.98) | <0.01 |
| Cardiac | Age | 2.19 | (2.1, 2.29) | <0.001 | 2.1 | (1.98, 2.21) | <0.001 | 2.11 | (2, 2.23) | <0.001 |
|  | Weight | 0.67 | (0.63, 0.71) | <0.001 | 0.71 | (0.66, 0.77) | <0.001 | 0.7 | (0.65, 0.75) | <0.001 |
|  | Age * Weight | - | - | - | 0.92 | (0.87, 0.97) | <0.01 | 0.93 | (0.88, 0.99) | 0.02 |
| Liver/Pancreas | Age | 2.12 | (2, 2.24) | <0.001 | 2.12 | (1.99, 2.26) | <0.001 | 2.12 | (1.98, 2.26) | <0.001 |
|  | Weight | 0.75 | (0.69, 0.81) | <0.001 | 0.75 | (0.68, 0.82) | <0.001 | 0.75 | (0.68, 0.82) | <0.001 |
|  | Age * Weight | - | - | - | 1.01 | (0.94, 1.08) | 0.85 | 1.02 | (0.95, 1.09) | 0.67 |
| Respiratory | Age | 2.01 | (1.89, 2.14) | <0.001 | 2.04 | (1.9, 2.2) | <0.001 | 2.06 | (1.91, 2.21) | <0.001 |
|  | Weight | 0.8 | (0.74, 0.87) | <0.001 | 0.78 | (0.71, 0.86) | <0.001 | 0.76 | (0.69, 0.84) | <0.001 |
|  | Age * Weight | - | - | - | 1.04 | (0.96, 1.13) | 0.32 | 1.06 | (0.98, 1.15) | 0.13 |

*Note:*
Age (years) and Weight (kg) are standardized by subtracting their means 7, 23 and dividing by their standard deviations 4, 13.
<sup>a</sup> A model with the main effects of age and weight.
<sup>b</sup> A model with the main effects of age, weight, and the interaction.
<sup>c</sup> A model with the main effects of variables in Model 2 plus adjusted for sex, purebred/mixed breed, and geographic region.

To understand the trend of disease history across age and size, we modeled lifetime prevalence of each disease category in three ways: 1) as a function of age and weight, 2) as a function of age, weight, and the age by weight interaction, and 3) as a function of age, weight, and their interaction, and adjusted for the dog’s sex, purebred/mixed breed, and census division. We used continuous age and weight and they were standardized by subtracting their means and dividing by their standard deviations. Dog’s sex was classified into four levels, neutered/intact male, or spayed/intact female, using both responses to the sex of the dog and whether the dog has been spayed or neutered. For census division, we used the reported state of address where the dog resides and divided them into the nine census divisions that are adopted by the United States Census Bureau (12).

Poisson regression with robust standard error estimates (13) was used to estimate prevalence ratios (PR) and construct 95% confidence intervals (CI) of the PRs. The Wald tests were to test whether lifetime prevalence increased with age, and whether the size of the dog influenced either the prevalence of a reported condition in the disease category or its pattern across age. Based on the estimated PRs with increasing weight from the unadjusted model without the interaction term, we separated the result tables and figures into two sections: one for disease categories more common in larger dogs and the other for disease categories less or equally common in larger dogs. In addition, we graphically illustrated the predicted prevalence of each disease category as a function of continuous age for each size class. These estimates were predicted using Poisson regression models fit with continuous age, categorical size, and the interaction of those two as predictors. Though fitted models included dogs aged 15+ years, we produced predictions for age from 0 to 15 as dogs older than 15 were very rare. Given the large sample size of the study and continuous nature of the exposures we used p<0.01 to denote statistical significance. For interactions with p<0.01 we also visually inspected the figures for qualitative evidence of important interaction.

All statistical analyses were performed in R 4.1.2 (14). The Dog Aging Project is an open data project. These data are available to the general public at dogagingproject.org/open_data_access (15).

## Results

There were 27,541 dog owners that completed the Health and Life Experience Survey and are included in this report. The dogs ranged in age from puppies to very senior dogs, with a median age of 7 years (IQR 4 to 11). The dogs were equally distributed by sex (50% male) and purebred versus mixed breed status (49% purebred). The respondents were distributed across the US, with the following percentages according to census divisions in decreasing order of representation: Pacific 24%, South Atlantic 18%, East North Central 15%, Mountain 10%, Mid-Atlantic 9%, West South Central 8%, New England 6%, West North Central 6%, East South Central 3%. The dogs most commonly resided in suburban locations (62%), with 17% residing in urban locations and 21% in rural locations. We now provide results for each disease category individually, with the categories that were positively associated with dog size discussed first, followed by those that were either negatively or not associated with dog size.

### Skin Conditions

A total of 7915 (28.7%) dog owners reported that their dogs had a history of conditions within the skin disease category (Table 1). The most commonly reported specific skin problems were seasonal allergies (n=1878) followed by pruritus (n=1164) and sebaceous cysts (n=943) (Supplementary Table 1). The proportion of dogs with a reported history of skin conditions increased steadily with age, ranging from 7% in puppies to 37% in senior dogs (Table 1). History of skin disease was reported less for smaller vs. larger dogs, ranging from 26% in dogs <10kg to 33% in dogs over 40kg (Table 2). In the Poisson regression model with the main effects of age and weight, each SD increment of age (4 years) was associated with a 29% greater relative prevalence of a reported history of skin conditions (PR 1.29, 95% CI 1.27-1.31, p<0.001). Prevalence also increased with the owner-reported weight of the dog, with each SD increment (13 lbs) associated with 12% higher relative prevalence of skin condition history (PR 1.12, 95% CI 1.10-1.14, p<0.001). There was no significant interaction between age and weight, which was in alignment with the results in Figure 1 that older dogs were reported to have skin conditions more often by a similar relative margin across different size classes. In Figure 1 we illustrate these patterns using a categorical representation of dog size. We see strongly increasing trends over age that are similar across size classes. Toy dogs have the lowest rates across age, and a history of skin conditions is reported progressively more often for each successively larger size category regardless of age at the time of the survey. Adjustment for sex, breed and geographic region did not have any notable impact on the associations, and this was true for all disease categories we studied.

**Figure 1:**
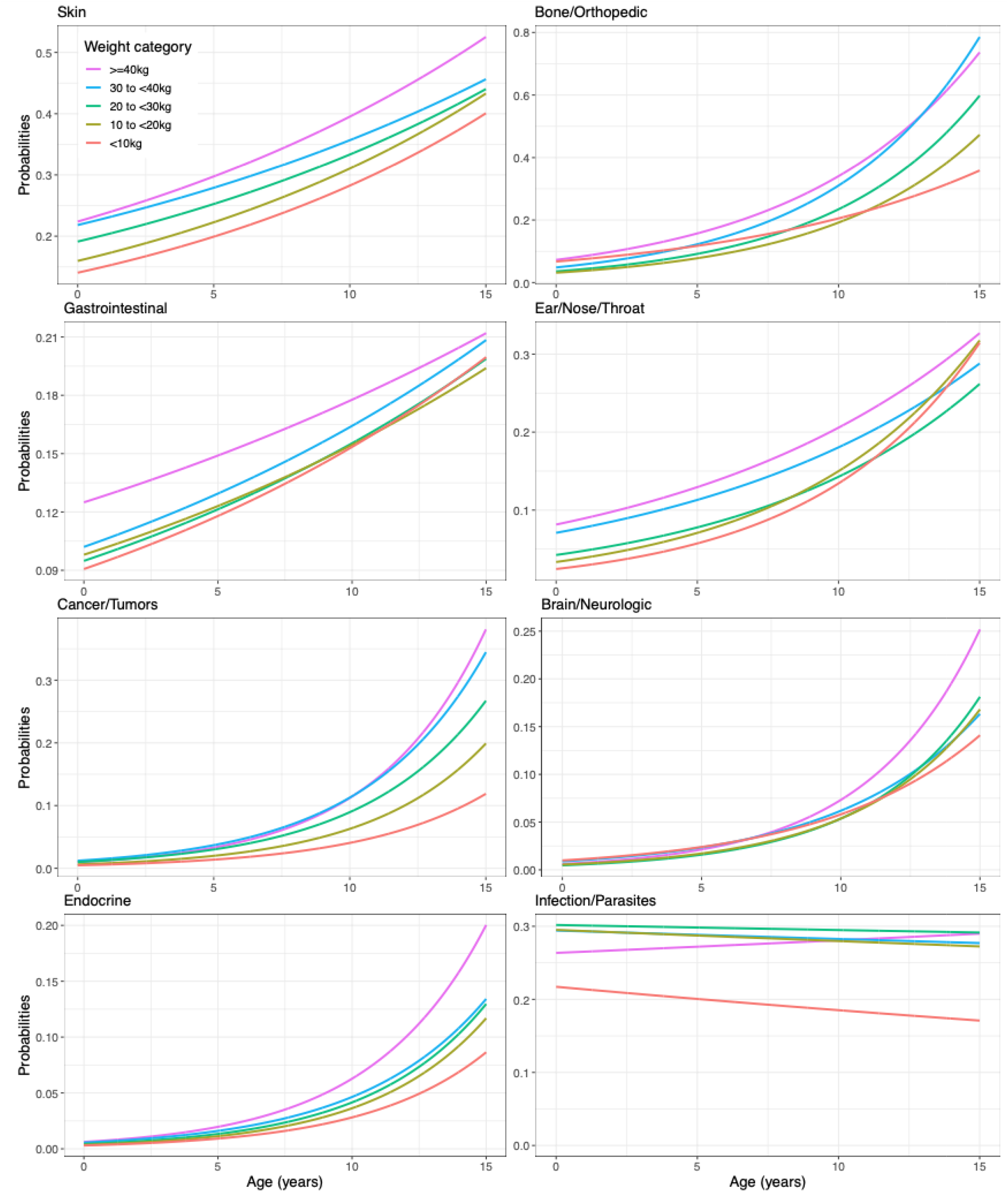
Results from Model 2 with Continuous Age by Weight Category (Part 1: Conditions More Common in Larger Dogs)

### Orthopedic Conditions

A total of 5287 (19.2%) dog owners reported that their dogs had a history of conditions within the orthopedic disease category (Table 1). The three most commonly reported specific orthopedic problems were osteoarthritis (n=1777) followed by cruciate ligament rupture (n=982) and patellar luxation (n=693) (Supplementary Table 1). Unsurprisingly, the proportion of dogs reported to have a history of orthopedic conditions increased sharply with age (from 1% in puppies to 39% in the oldest dog group, Table 1) and slightly with size (from 19% in the <10kg dogs to 23% in the dogs over 40kg, Table 2). On average, the prevalence of a reported history of orthopedic conditions doubled with every SD increment in age (PR 2.00, 95% CI 1.95-2.04, p<0.001), but as indicated by the interaction term, the prevalence across age increased much more sharply as the size of the dog increased. This is illustrated in Figure 1, where for puppies and adolescents (<3 years) there is very little difference in prevalence by size class, but for older adult to senior (7+ years) the larger dogs have a much greater prevalence of a history of orthopedic conditions.

### Gastrointestinal (GI) Conditions

A total of 3914 (14.2%) dog owners reported that their dogs had a history of conditions within the GI disease category (Table 1). The most commonly reported specific GI conditions were chronic or recurrent diarrhea (n=786) followed by anal sac impaction (n=720) and foreign body ingestion or blockage (n=685) (Supplementary Table 1). The reported proportion of dogs with a history of GI conditions increased with age (from 7% in puppies to 18% among senior dogs) but only slightly with size (14% in dogs <10kg to 16% in dogs over 40kg). Each SD increment in dog age was associated with a 22% higher relative prevalence (PR 1.22, 95% CI 1.19-1.26, p<0.001), while each SD increment in dog weight was associated with 4% greater relative prevalence (PR 1.04, 95% CI 1.01-1.07, p=0.01). In Figure 1 we see that all size groups increase steadily with age. Dogs <30kg are all similar, dogs between 30-40kg have somewhat higher prevalence, and dogs over 40kg have notably higher prevalence across all ages.

### Ear, Nose, and Throat (ENT) Conditions

A total of 3569 (13%) dog owners reported that their dogs had a history of conditions within the ENT disease category (Table 1). The three most commonly reported specific ENT conditions were chronic or recurrent ear infections (n=1998) followed by hearing loss (n=653) and acquired deafness (n=405) (Supplementary Table 1). Reported history of ENT conditions increased from 3% among puppies to 25% among senior dogs. Trends across size were slight, ranging from 13% in the smallest dogs (<10kg) to 16% in the largest dogs (>=40kg). Controlling for the weight of the dog, on average the relative prevalence of ENT conditions was 74% higher for each SD increment in age (PR 1.76, 95% CI 1.70-1.82, p<0.001). However, there was a significant interaction between size and weight. Specifically, for larger dogs the trend across age was flatter. Puppies to young adults (<7 years) had higher prevalence of ENT history for larger dogs, but for seniors (11+ years), small dogs (<20 kg) had caught up to the largest dogs. This is illustrated in Figure 1.

### Cancer/Tumors

A total of 1751 (6.4%) dog owners reported that their dogs had a history of conditions within the cancer/tumor category (Table 1). The most commonly reported specific cancer sites were skin of trunk/body/head (n=502) followed by muscle or other soft tissue (n=376) and skin of limb/foot (n=254). The most commonly known types of cancer were mast cell tumor (n=349) and lipoma (n=217). However, many owners were unsure of types of cancer (n=499) (Supplementary Table 1). The proportion of dogs for whom a history of cancer was reported increased sharply with age, ranging from <1% in puppies and adolescent dogs, 2% in young adult dogs, 8% in older adult dogs, and 15% in senior dogs. Prevalence by size ranged from 4% in dogs <10kg, to between 6-8% among dogs over 10kg. The increasing pattern across age was much more pronounced in larger dogs. For dogs of the same size, each SD increment in dog age was associated with a two-and-a-half-fold increase in the prevalence of cancer history (PR 2.56, 95% CI 2.45-2.67, p<0.001). For each additional SD increment in size, the age trend increased by 7%. In Figure 1 we see this illustrated graphically, where for puppies and adolescents (<3 years), prevalence is low for all sizes, but for seniors (11+ years) the prevalence has risen rapidly, especially for dogs over 30kg.

### Neurologic Conditions

A total of 1324 (4.8%) dog owners reported that their dogs had a history of conditions within the neurologic disease category (Table 1). The three most commonly reported specific neurologic conditions were seizures (n=606) followed by dementia/senility (n=153) and vestibular disease (n=147) (Supplementary Table 1). The proportion of dogs with a history of neurologic conditions increased from <1% in puppies and adolescent dogs to 12% in senior dogs. On average there was no apparent trend across sizes, though the largest dogs had a much steeper increasing pattern across age than dogs <40kg. Before older adulthood the prevalence of neurologic conditions was low, and similar across size classes. In older adult and senior dogs the prevalence increased in a similar way for dogs <40kg, and more steeply for dogs over 40kg.

### Endocrine Conditions

A total of 913 (3.3%) dog owners reported that their dogs had a history of conditions within the endocrine disease category (Table 1). The three most commonly reported specific endocrine conditions were hypothyroidism (n=520), followed by Cushing’s disease (n=175) and diabetes mellitus (n=83) (Supplementary Table 1). The prevalence of a reported history of endocrine conditions was <1% through young adulthood, 4% for older adult dogs, and 8% for senior dogs. On average there was not a strong size pattern with prevalence ranging between 3-4%. However, controlling for age the influence of size on prevalence was apparent (PR 1.27, 95% CI 1.19, 1.35, p<0.001). The increasing pattern of endocrine condition history across age was similar for all size classes. These patterns are illustrated in Figure 1, where the prevalence of reported history of endocrine conditions increases over time at a similar rate for each size class, but the larger the dog the higher the estimated prevalence curve.

### Infectious Diseases

A total of 7339 (26.6%) dog owners reported that their dogs had a history of conditions within the infectious disease category (Table 1). The three most commonly reported specific infectious conditions were *Giardia* (n=1958) followed by *Bordetella* and/or parainfluenza (kennel cough) (n=1277) and tapeworms (n=816) (Supplementary Table 1). The proportion of dogs with a reported history of infectious diseases followed a unique pattern relative to all other conditions studied. The prevalence among puppies was 25%, and this did not increase across other age groups ranging between 25-28% with no particular trend. The smallest dogs <10kg had the lowest reported history of infectious diseases at 19%, but all other size classes were similar to each other, with 27-30% prevalence. These patterns are most clearly illustrated in Figure 1, where the pattern across age groups is flat, and the curve for the smallest dogs is lowest.

### Ocular Conditions

A total of 3625 (13.2%) dog owners reported that their dogs had a history of conditions within the ocular disease category (Table 1). The three most commonly reported specific ocular conditions were adult-onset cataracts (n=1114) followed by conjunctivitis (n=759) and corneal ulcers (n=321) (Supplementary Table 1). A reported history of ocular conditions increased steadily with age group, from 5% in puppies to 28% among senior dogs. In contrast, the proportion with a positive history was lower among larger dogs, ranging from 17% in the smallest dogs to just 10% in the dogs over 40kg. This was reflected in the models where the significant interaction between age and weight is seen as a steeper slope in smaller dogs. Graphically we can see in Figure 2 that below adulthood the size categories have roughly the same prevalence but then the smaller dogs increase much more quickly in reported history of ocular conditions.

**Figure 2:**
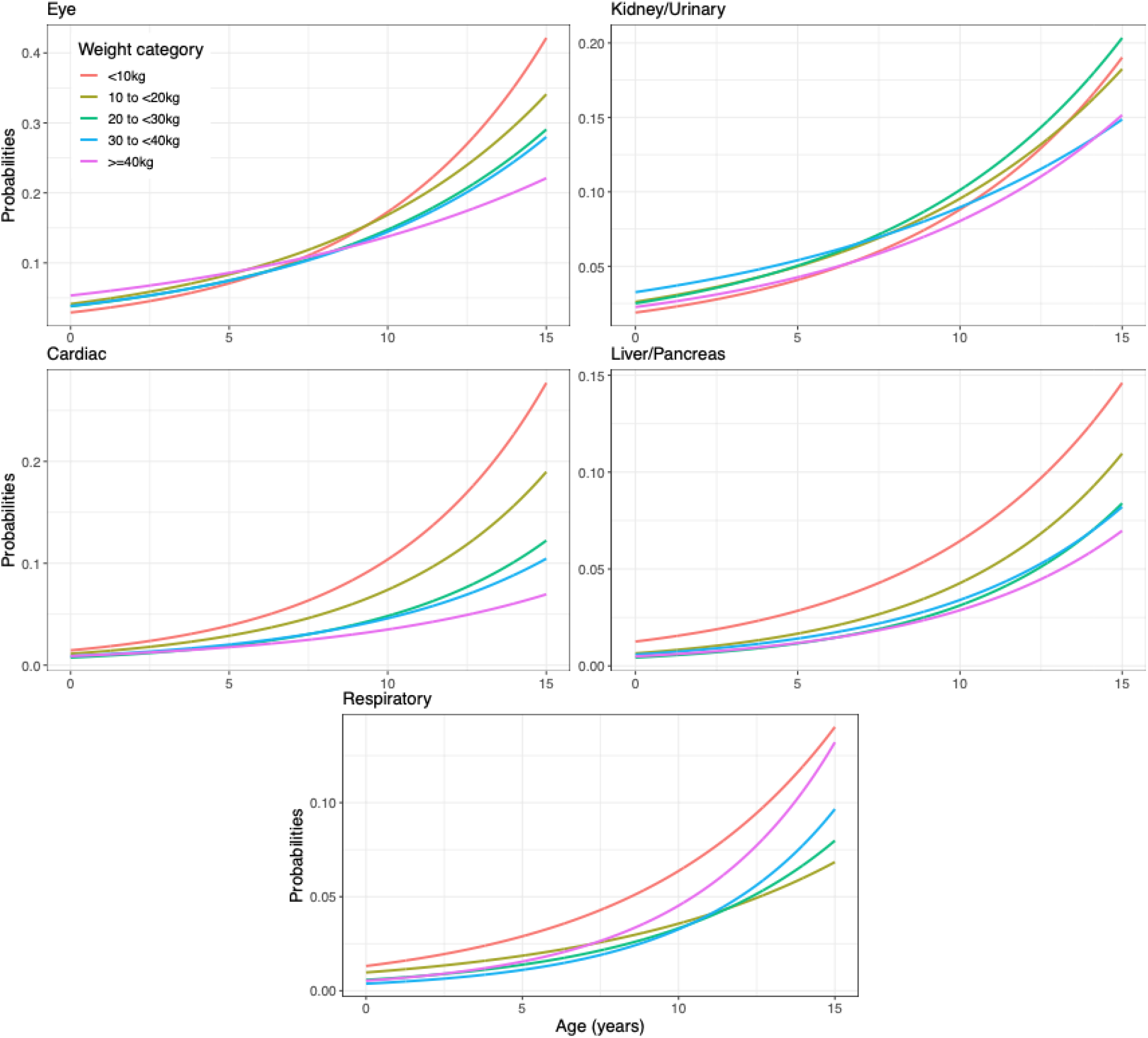
Results from Model 2 with Continuous Age by Weight Category (Part 2: Conditions Less or Equally Common in Larger Dogs)

### Kidney or Urinary Conditions

A total of 2122 (7.7%) dog owners reported that their dogs had a history of conditions within the kidney or urinary disease category (Table 1). The three most commonly reported specific kidney or urinary conditions were chronic or recurrent urinary tract infection (n=813) followed by urinary incontinence (n=636) and urinary crystals or stones in bladder or urethra (n=423) (Supplementary Table 1). The proportion of dogs with reported history of kidney or urinary conditions increased steadily with age, from 2% in puppies to 15% in senior dogs. Across size categories differences were not dramatic, but the prevalence was highest in the smallest dogs at 9% in dogs <10kg and lowest in largest dogs at 6% in dogs >=40kg. Differences between size classes were most apparent in older adults and seniors, as evidenced by the significant age by weight interaction, and the graphical representation in Figure 2.

### Cardiac conditions

A total of 1567 (5.7%) dog owners reported that their dogs had a history of conditions within the cardiac disease category (Table 1). The three most commonly reported specific cardiac conditions were murmur (n=1156) followed by valve disease (n=139) and congestive heart failure (n=137) (Supplementary Table 1). The proportion of dogs with a reported history of cardiac conditions exhibited strong trends across both age and dog size. Among puppies and adolescents the proportion was <1%, increasing to 14% among senior dogs. Dog weight was inversely associated, with dogs <10kg reporting an 11% prevalence of a history of cardiac conditions, declining to only 2% among the largest dogs over 40kg. These trends were highly significant, with each SD increment in age associated with over a doubling of prevalence (PR 2.19, 95% CI 2.1-2.29, p<0.001), and each SD increment in size reducing prevalence by a third (PR 0.67, 95% CI 0.63-0.71, p<0.001). Additionally, we observed a significant interaction where the prevalence was not only higher for smaller dogs but was associated with a significantly steeper increase in prevalence across age groups.

### Liver or Pancreas Conditions

A total of 970 (3.5%) dog-owners reported that their dogs had a history of conditions within the liver or pancreas disease category (Table 1). The three most commonly reported specific liver or pancreas conditions were pancreatitis (n=587) followed by chronic inflammatory liver disease (n=86) and exocrine pancreatic insufficiency (n=35) (Supplementary Table 1). A reported history of liver or pancreas conditions was more common among older dogs ranging from <1% in puppies and adolescent dogs to 8% in senior dogs. In contrast the proportion was lower among larger dogs, ranging from 6% in the smallest size category down to 2% among larger dogs. Each SD increment in age more than doubled the prevalence of a reported history of liver/pancreas conditions (PR 2.12, 95% CI 2.00 to 2.24, p<001), while each SD increment in size reduced the prevalence by approximately one quarter (PR 0.75, 95% CI 0.69-0.81, p<0.001). The smallest dog category (<10kg) in particular had a notably higher prevalence of liver condition history.

### Respiratory Conditions

A total of 950 (3.4%) dog owners reported that their dogs had a history of conditions within the respiratory disease category (Table 1). The three most commonly reported specific respiratory conditions were chronic or recurrent cough (n=236) followed by tracheal collapse (n=169) and pneumonia (n=163) (Supplementary Table 1). The proportion of dogs with a reported history of respiratory conditions went up with increased age, from <1% in puppies to 8% in senior dogs. The proportion decreased with increasing size, from 6% in the smallest dogs to 3% in the largest dogs. For every 1 SD increment in age the prevalence of a reported history of respiratory conditions doubled (PR 2.01, 95% CI 1.89-2.14, p<0.001), while for every 1 SD increment in weight the relative prevalence was 20% lower (PR 0.80, 95% CI 0.74-0.87, p<0.001). These patterns are illustrated in the final panel of Figure 2.

## Discussion

We are interested in identifying potential patterns of disease contributing to decreased average lifespan with increasing size in pet dogs. As found in other retrospective studies of the domestic dog (1,10,16–20), we found that owner-reported diagnoses (grouped by organ system or pathophysiologic process) disproportionately affect dogs of different sizes. Some conditions were reported less commonly for larger dogs and as such are unlikely to be an important contributor to their shorter lifespan. These included ocular, cardiac, liver/pancreas, and respiratory conditions. The proportion reporting a history of urinary conditions did not vary by weight. Many conditions were reported more commonly with increasing weight category, including skin, orthopedic, gastrointestinal, ear/nose/throat, cancer, neurologic, and endocrine conditions. The infectious disease category showed a distinct pattern that the smallest dogs (<10kg) had much lower prevalence than the other categories and there were no increasing patterns across age. With the exception of infectious diseases, the proportion of dogs with a reported history of each disease category increased with increasing age group as expected. Effects of size included differences in relative prevalence of reporting a history of disease as well as variable patterns of increase in prevalence across age categories.

Higher growth rates in larger dogs have been implicated in increased oxidative damage during early life that may predispose to certain diseases such as skin diseases, orthopedic conditions, cancer, and cardiac disorders (21). Skin is a common site for evidence of oxidative damage; increased production of free radicals including reactive oxygen species (ROS) can overwhelm the body’s mechanisms for reducing their damaging effects and manifest as skin lesions and other pathology (22). In mouse models, increased oxidative stress induced by exposure to diisodecyl phthalate (DIDP) exacerbated symptoms of allergic dermatitis (23). A retrospective study of 721 dogs by Dreschel (24) measuring associations between behavioral health and physical health found that dogs with severe fear and separation anxiety disorders had both significantly higher prevalence and severity of skin diseases as well as significantly shorter lifespan (24). The author suggests that these associations may be due to physiological stress responses in dogs with behavioral disorders and resulting changes in hormone regulation, immunity, and disease risk.

Faster growth rates and larger body size have also been associated with increased risk for orthopedic diseases (6). Growth patterns in larger dog breeds involve rapid weight gain throughout the period of skeletal development and maturity, which has been linked to increased risk of developmental musculoskeletal and orthopedic diseases including hip dysplasia, osteoarthritis, and osteochondrosis (6,25,26). These conditions have been implicated in reducing longevity in Labrador Retrievers (27). In our cohort, osteoarthritis (OA), was spread similarly among dogs with body weight greater than ten kilograms in our cohort, the morbidity associated with OA may vary. Orthopedic pain from any of these diagnoses can significantly reduce quality of life (27). Poor QOL has been identified as one of the most influential factors for owners choosing to euthanize their dogs and thereby shorten their lifespan to prevent pain and distress (28). Most medications commonly used to alleviate pain from OA are dosed by weight and are thus more likely to be cost-prohibitive for owners with larger dogs; such factors may also lead to differential effects of OA by size on QOL and decisions about euthanasia. Additionally, mobility problems due to orthopedic disease are more easily accommodated in dogs of smaller size, as owners are more likely to be capable of lifting or carrying smaller dogs and/or assisting their movement with slings, harnesses, and other devices.

Across all body sizes, several retrospective studies of large cohorts of companion dogs in North America and Europe have consistently found that cancer (1,17,19,20,28,29) is one the most common causes of mortality in pet dogs. Multiple studies that have considered body size in relation to causes of morbidity and mortality have reported that cancer diagnoses are more common in larger dogs than smaller dogs (1,30–32). Several theories about the contributing factors to reduced longevity in larger dogs are consistent with increased risk of cancer in larger dogs including correlations between increased serum IGF-1 levels in larger dogs and downstream effects on growth, oxidative damage, and cancer risk (6,30,33,34). Artificial selection to produce the extreme variation in growth rates between breeds has resulted in wide variation in serum IGF-1 levels before and after skeletal maturity. For example, Great Danes gained weight at 17 times the rate of miniature poodles in a study following the first 21 weeks of life (35). Correspondingly higher levels of mean basal IGF-1 detected in the plasma of Great Danes compared to miniature poodles from age 13 weeks to 27 weeks in a related study suggested that Great Danes may enter a period of physiologic gigantism postnatally (36). Plasma levels of IGF-1 remain higher in larger breed dogs compared to smaller breed dogs past maturity, and this inverse correlation between IGF-1 levels and longevity is consistent with patterns found in many other species (6).

Studies investigating the relationship between average breed growth rates and prevalence of life-limiting disorders linked to oxidative damage have revealed correlations between breed size and risk of developing cardiac pathology. Increased risk of cardiac disease and associated mortality has been reported at both extremes of the body size spectrum in dogs, though the prevalence of specific cardiac pathophysiologies varies widely from small dogs to giant breed dogs (1,19,20). Artificial selection for extreme size (both small and giant) may have contributed to the distribution of cardiac disease risk in dogs, as many genes associated with body size in dogs also contribute to cardiac development and structure (37). Telomere length in peripheral blood mononuclear cells was positively correlated with lifespan and inversely correlated with risk of mortality due to cardiac disease in a study of 15 breeds of dog, with the shortest age-adjusted telomere length in the Great Dane (38). In our cohort, reported prevalence of cardiac disease increases more rapidly with age in smaller dogs. Contribution of cardiac disease to mortality risk in our cohort is unknown.

Reported prevalence of diagnoses in the gastrointestinal category was more common as size increased in our cohort. The most commonly reported diagnoses in this category (chronic or recurrent diarrhea, anal sac impactions, foreign body ingestion or blockage, and food or medicine allergies) can vary widely in severity, and therefore potential effects on longevity are difficult to predict. Increased risk of mortality due to gastrointestinal disease with increasing body size has been reported by Fleming et al. (1).

In our cohort, larger dogs were also more likely to have diagnoses in the neurological category. Of these diagnoses “seizures (including epilepsy)” was the most commonly reported specific condition, and was distributed fairly evenly across body sizes. Fleming, et al. (1) found that larger dogs were relatively spared from death due to neurological diseases. Our results are not inconsistent with this finding, as many dogs with seizures do not die as a direct result of their neurological disease; the majority of dogs with epilepsy (the most common cause of recurrent seizures in dogs) can be treated successfully with conventional drugs (39) and, similarly, the majority of companion dogs with epilepsy die of a cause not directly related to epilepsy (40). Additionally, neurological diseases associated with greater mortality risk compared to epilepsy are more common in smaller dogs (e.g., intervertebral disc disease), and tend to present later in life than epilepsy, such that neurological disease in smaller dogs would be less frequent in our cohort than in cohorts examined retrospectively for causes of mortality.

Previously, Fleming et al. (1) reported that larger breeds are spared from death by endocrine disease. In our cohort, the reported prevalence of a diagnosis in the endocrine category was more likely in larger dogs. This contrast between reported mortality versus prevalence may be driven almost entirely by the most commonly reported specific endocrine disorder, hypothyroidism. Hypothyroidism was more prevalent in larger dogs, and this diagnosis alone was responsible for most of the trend by size for the endocrine category. While untreated hypothyroidism can cause significant morbidity and even mortality, it is relatively easy and inexpensive to treat in dogs (41,42). In contrast, Cushing’s disease and diabetes mellitus, two frequently diagnosed diseases that are more prevalent in smaller dogs in our cohort, carry greater mortality risk and financial burden for the owner (43–46). It is noteworthy that Fleming et al. (1) used data from dogs seen in referral institutions while the dogs in our study were recruited directly from the community. It is therefore likely that referral bias (47) could also explain the difference in these results, as treatment of hypothyroidism is usually straightforward and does not require referral (48).

For the ENT category, the most commonly reported diagnosis, ear infection, was drastically skewed toward larger dogs. Ear infections in dogs are commonly associated with allergic skin disease and food allergies and thus increased prevalence in larger dogs would be consistent with the aforementioned theories about increased oxidative damage and skin manifestations. While ear infections can sometimes be easily treated, chronic/recurrent ear infections are also common and can drastically affect quality of life (49–52). Recurrent ear infections can be expensive to monitor and treat, with surgery to remove the entire ear canal(s) as the only curative option for some dogs. Because significant financial resources may be necessary to avoid negative impacts on quality of life due to ear infections, cost may be a factor guiding euthanasia decisions and thus lifespan (52,53). The next most frequent diagnoses in the ENT category were hearing loss and deafness, which both have frequencies skewed toward smaller dogs. In general, pet dogs do not receive treatment for hearing loss or deafness, nor do they receive adaptive devices such as hearing aids and thus financial limitations of the owner do not tend to change the prognosis for the patient with chronic ear infections. Hearing loss is typically not associated with pain, and dogs are often capable of maintaining a good quality of life despite partial or complete deafness.

Our study has several strengths and limitations that should be noted. Strengths include the large sample size of this study, which allows us to estimate patterns accurately across the whole age and size spectrum. Additionally, we have a very diverse sample of dogs distributed across the entire United States. Since the sample is not veterinary-hospital or clinic-based it may be more representative of the general population of dogs. Conversely, while our observations can suggest which conditions manifest differently across age and size, they do not prove any causal relationships due to the cross-sectional nature of the analysis. Over time, longitudinal data will be collected on these dogs, and we will be able to examine disease incidence. In addition, recall bias may occur when owners fill out the survey. It is possible that owners may not remember past events at all or incorrectly at the time of the survey. Future studies will compare HLES data to Veterinary Electronic Medical Records data to measure the accuracy of owner’s reported diagnoses and confirm the trends seen in HLES. Prevalence of a history of conditions was expected to go up regardless of an increasing age-specific prevalence because it was calculated in a cumulative way. Finally, the sample is not random but self-selected so that data are subject to self-selection bias. Owners who are more exposed to this survey and who tend to participate in surveys are more likely to nominate their dogs and complete the survey, possibly leading to a biased sample. The Dog Aging Project endeavors to create a representative sample of the companion canine population. However, computer access is a necessary condition for survey completion, as well as the ability to complete the surveys in English.

In this study we have quantified the reported prevalence of a history of conditions within several different disease categories as a function of dog age and size, with and without adjustment for sex, geographic location, and pure versus mixed breed status. These results provide insights into the disease categories that may contribute to reduced lifespan in larger dogs and suggest multiple further avenues for further exploration. More focused efforts to look at individual conditions within categories may yield additional insights. Within and across categories, the co-occurrence of different disease subtypes may also be an important factor to evaluate. Of course, as prospective data become available the longitudinal associations of these conditions with subsequent morbidity and mortality will be evaluated.

## Acknowledgments

This research is based on publicly available data collected by the Dog Aging Project, which is supported by U19 grant AG057377 from the National Institute on Aging, a part of the National Institutes of Health, and by additional grants and private donations. These data are housed on the Terra platform at the Broad Institute of MIT and Harvard. The content is solely the responsibility of the authors and does not necessarily represent the official views of the National Institutes of Health.

The Dog Aging Project thanks study participants, their dogs, and community veterinarians for their important contributions.

