## Supplemental Tables for "Dog Size and Patterns of Disease History Across the Canine Age Spectrum: Results from the Dog Aging Project": supplementary-tables.html

condition-by-weight.utf8


Table 1: Proportion with Skin Disorder History by Weight Category

|  | Weight Category | | | | |  |
| --- | --- | --- | --- | --- | --- | --- |
| Skin | <10kg (N=1628) | 10 to <20 (N=1531) | 20 to <30 (N=2370) | 30 to <40 (N=1621) | >=40 (N=766) | Total (N=7916) |
| Seasonal allergies | 382 (23%) | 368 (24%) | 538 (23%) | 404 (25%) | 186 (24%) | 1878 (24%) |
| Pruritis (itchy skin) | 272 (17%) | 231 (15%) | 337 (14%) | 241 (15%) | 83 (11%) | 1164 (15%) |
| Sebaceous cysts | 221 (14%) | 189 (12%) | 276 (12%) | 185 (11%) | 72 (9%) | 943 (12%) |
| Food or medicine allergies that affect the skin | 146 (9%) | 152 (10%) | 247 (10%) | 182 (11%) | 115 (15%) | 842 (11%) |
| Chronic or recurrent hot spots | 131 (8%) | 130 (8%) | 216 (9%) | 215 (13%) | 136 (18%) | 828 (10%) |
| Fleas | 228 (14%) | 184 (12%) | 224 (9%) | 125 (8%) | 49 (6%) | 810 (10%) |
| Other | 174 (11%) | 136 (9%) | 242 (10%) | 163 (10%) | 69 (9%) | 784 (10%) |
| Atopic dermatitis (atopy) | 164 (10%) | 135 (9%) | 243 (10%) | 161 (10%) | 71 (9%) | 774 (10%) |
| Ticks | 115 (7%) | 153 (10%) | 239 (10%) | 128 (8%) | 57 (7%) | 692 (9%) |
| Chronic or recurrent skin infections | 96 (6%) | 87 (6%) | 122 (5%) | 120 (7%) | 66 (9%) | 491 (6%) |
| Flea allergy dermatitis | 120 (7%) | 80 (5%) | 114 (5%) | 67 (4%) | 26 (3%) | 407 (5%) |
| Contact dermatitis | 71 (4%) | 71 (5%) | 105 (4%) | 110 (7%) | 38 (5%) | 395 (5%) |
| Alopecia (hair loss) | 92 (6%) | 63 (4%) | 113 (5%) | 69 (4%) | 19 (2%) | 356 (4%) |
| Non specific dermatosis | 56 (3%) | 61 (4%) | 110 (5%) | 73 (5%) | 40 (5%) | 340 (4%) |
| Lick granuloma | 32 (2%) | 53 (3%) | 82 (3%) | 80 (5%) | 39 (5%) | 286 (4%) |
| Pyoderma or bacterial dermatitis | 46 (3%) | 52 (3%) | 75 (3%) | 56 (3%) | 30 (4%) | 259 (3%) |
| Systemic demodectic mange | 17 (1%) | 39 (3%) | 55 (2%) | 32 (2%) | 12 (2%) | 155 (2%) |
| Sarcoptic mange | 22 (1%) | 28 (2%) | 47 (2%) | 24 (1%) | 10 (1%) | 131 (2%) |
| Seborrhea or seborrheic dermatitis (greasy skin) | 24 (1%) | 12 (<1%) | 17 (<1%) | 14 (<1%) | 4 (<1%) | 71 (<1%) |
| Pododermatitis | 7 (<1%) | 10 (<1%) | 16 (<1%) | 13 (<1%) | 2 (<1%) | 48 (<1%) |
| Discoid lupus erythematosus (DLE) | 3 (<1%) | 3 (<1%) | 6 (<1%) | 5 (<1%) | 3 (<1%) | 20 (<1%) |
| Ichthyosis | 1 (<1%) | 0 (<1%) | 8 (<1%) | 7 (<1%) | 0 (<1%) | 16 (<1%) |
| Sebaceous adenitis | 3 (<1%) | 4 (<1%) | 5 (<1%) | 2 (<1%) | 0 (<1%) | 14 (<1%) |
| Pemphigus foliaceus (PF) | 2 (<1%) | 2 (<1%) | 3 (<1%) | 3 (<1%) | 1 (<1%) | 11 (<1%) |
| Systemic lupus erythematosus (SLE) | 0 (<1%) | 0 (<1%) | 3 (<1%) | 1 (<1%) | 0 (<1%) | 4 (<1%) |
| Panepidermal pustular pemphigus (PPP) | 0 (<1%) | 1 (<1%) | 1 (<1%) | 0 (<1%) | 0 (<1%) | 2 (<1%) |
| Pemphigus erythematosus (PE) | 0 (<1%) | 0 (<1%) | 0 (<1%) | 1 (<1%) | 0 (<1%) | 1 (<1%) |
| Pemphigus vulgaris (PV) | 1 (<1%) | 0 (<1%) | 0 (<1%) | 0 (<1%) | 0 (<1%) | 1 (<1%) |
| Paraneoplastic pemphigus (PNP) | 0 (<1%) | 0 (<1%) | 0 (<1%) | 0 (<1%) | 0 (<1%) | 0 (<1%) |
| Polymyositis | 0 (<1%) | 0 (<1%) | 0 (<1%) | 0 (<1%) | 0 (<1%) | 0 (<1%) |

Table 2: Proportion with Infectious or Parasitic Disease History by Weight Category

|  | Weight Category | | | | |  |
| --- | --- | --- | --- | --- | --- | --- |
| Infectious | <10kg (N=1186) | 10 to <20 (N=1593) | 20 to <30 (N=2441) | 30 to <40 (N=1478) | >=40 (N=641) | Total (N=7339) |
| Giardia | 330 (28%) | 450 (28%) | 638 (26%) | 373 (25%) | 167 (26%) | 1958 (27%) |
| Bordetella and or parainfluenza (kennel cough) | 209 (18%) | 263 (17%) | 438 (18%) | 257 (17%) | 110 (17%) | 1277 (17%) |
| Tapeworms | 136 (11%) | 171 (11%) | 276 (11%) | 164 (11%) | 69 (11%) | 816 (11%) |
| Roundworms | 107 (9%) | 167 (10%) | 263 (11%) | 147 (10%) | 94 (15%) | 778 (11%) |
| Lyme disease | 93 (8%) | 153 (10%) | 246 (10%) | 160 (11%) | 58 (9%) | 710 (10%) |
| Hookworms | 94 (8%) | 172 (11%) | 242 (10%) | 154 (10%) | 46 (7%) | 708 (10%) |
| Gastrointestinal parasites | 78 (7%) | 127 (8%) | 177 (7%) | 115 (8%) | 46 (7%) | 543 (7%) |
| Heartworm infection | 63 (5%) | 98 (6%) | 195 (8%) | 86 (6%) | 33 (5%) | 475 (6%) |
| Coccidia | 78 (7%) | 98 (6%) | 144 (6%) | 78 (5%) | 40 (6%) | 438 (6%) |
| Other | 51 (4%) | 88 (6%) | 110 (5%) | 60 (4%) | 29 (5%) | 338 (5%) |
| Anaplasmosis | 30 (3%) | 68 (4%) | 119 (5%) | 47 (3%) | 37 (6%) | 301 (4%) |
| Whipworms | 35 (3%) | 67 (4%) | 100 (4%) | 44 (3%) | 23 (4%) | 269 (4%) |
| Ehrlichiosis | 22 (2%) | 51 (3%) | 98 (4%) | 51 (3%) | 21 (3%) | 243 (3%) |
| Parvovirus | 40 (3%) | 45 (3%) | 70 (3%) | 59 (4%) | 22 (3%) | 236 (3%) |
| Dermatophytosis (ringworm) | 14 (1%) | 25 (2%) | 32 (1%) | 16 (1%) | 7 (1%) | 94 (1%) |
| Fever of unknown origin | 13 (1%) | 16 (1%) | 24 (<1%) | 11 (<1%) | 6 (<1%) | 70 (<1%) |
| Influenza | 10 (<1%) | 7 (<1%) | 11 (<1%) | 9 (<1%) | 7 (1%) | 44 (<1%) |
| Leptospirosis | 12 (1%) | 6 (<1%) | 12 (<1%) | 8 (<1%) | 5 (<1%) | 43 (<1%) |
| MRSA MRSP | 7 (<1%) | 3 (<1%) | 11 (<1%) | 15 (1%) | 6 (<1%) | 42 (<1%) |
| Coccidioidomycosis | 4 (<1%) | 4 (<1%) | 12 (<1%) | 9 (<1%) | 2 (<1%) | 31 (<1%) |
| Campylobacteriosis | 4 (<1%) | 5 (<1%) | 10 (<1%) | 3 (<1%) | 4 (<1%) | 26 (<1%) |
| Distemper | 2 (<1%) | 8 (<1%) | 6 (<1%) | 5 (<1%) | 1 (<1%) | 22 (<1%) |
| Babesiosis | 4 (<1%) | 2 (<1%) | 9 (<1%) | 5 (<1%) | 1 (<1%) | 21 (<1%) |
| Rocky Mountain Spotted Fever (RMSF) | 3 (<1%) | 2 (<1%) | 8 (<1%) | 0 (<1%) | 8 (1%) | 21 (<1%) |
| Granuloma | 0 (<1%) | 4 (<1%) | 6 (<1%) | 8 (<1%) | 2 (<1%) | 20 (<1%) |
| Salmon poisoning | 0 (<1%) | 3 (<1%) | 8 (<1%) | 6 (<1%) | 1 (<1%) | 18 (<1%) |
| Isospora | 3 (<1%) | 3 (<1%) | 6 (<1%) | 4 (<1%) | 1 (<1%) | 17 (<1%) |
| Blastomycosis | 0 (<1%) | 2 (<1%) | 4 (<1%) | 2 (<1%) | 4 (<1%) | 12 (<1%) |
| Aspergillosis | 0 (<1%) | 1 (<1%) | 6 (<1%) | 1 (<1%) | 0 (<1%) | 8 (<1%) |
| Cryptococcus | 0 (<1%) | 2 (<1%) | 4 (<1%) | 2 (<1%) | 0 (<1%) | 8 (<1%) |
| Mycobacterium | 0 (<1%) | 0 (<1%) | 4 (<1%) | 2 (<1%) | 0 (<1%) | 6 (<1%) |
| Salmonellosis | 0 (<1%) | 0 (<1%) | 0 (<1%) | 5 (<1%) | 1 (<1%) | 6 (<1%) |
| Toxoplasma | 0 (<1%) | 0 (<1%) | 3 (<1%) | 1 (<1%) | 0 (<1%) | 4 (<1%) |
| Brucellosis | 0 (<1%) | 0 (<1%) | 2 (<1%) | 1 (<1%) | 0 (<1%) | 3 (<1%) |
| Histoplasmosis | 1 (<1%) | 0 (<1%) | 0 (<1%) | 1 (<1%) | 0 (<1%) | 2 (<1%) |
| Leishmaniasis | 0 (<1%) | 0 (<1%) | 0 (<1%) | 1 (<1%) | 0 (<1%) | 1 (<1%) |
| Plague (Yersinia pestis) | 0 (<1%) | 0 (<1%) | 1 (<1%) | 0 (<1%) | 0 (<1%) | 1 (<1%) |
| Tularemia | 0 (<1%) | 0 (<1%) | 1 (<1%) | 0 (<1%) | 0 (<1%) | 1 (<1%) |
| Chagas disease (trypanosomiasis) | 0 (<1%) | 0 (<1%) | 0 (<1%) | 0 (<1%) | 0 (<1%) | 0 (<1%) |
| Hepatozoonosis | 0 (<1%) | 0 (<1%) | 0 (<1%) | 0 (<1%) | 0 (<1%) | 0 (<1%) |
| Pythium | 0 (<1%) | 0 (<1%) | 0 (<1%) | 0 (<1%) | 0 (<1%) | 0 (<1%) |

Table 3: Proportion with Orthopedic Disorder History by Weight Category

|  | Weight Category | | | | |  |
| --- | --- | --- | --- | --- | --- | --- |
| Orthopedic | <10kg (N=1187) | 10 to <20 (N=931) | 20 to <30 (N=1461) | 30 to <40 (N=1181) | >=40 (N=527) | Total (N=5287) |
| Osteoarthritis | 246 (21%) | 323 (35%) | 584 (40%) | 430 (36%) | 194 (37%) | 1777 (34%) |
| Cruciate ligament rupture | 104 (9%) | 146 (16%) | 334 (23%) | 267 (23%) | 131 (25%) | 982 (19%) |
| Patellar luxation | 492 (41%) | 104 (11%) | 56 (4%) | 30 (3%) | 11 (2%) | 693 (13%) |
| Hip dysplasia | 61 (5%) | 72 (8%) | 209 (14%) | 200 (17%) | 98 (19%) | 640 (12%) |
| Other | 112 (9%) | 122 (13%) | 156 (11%) | 112 (9%) | 50 (9%) | 552 (10%) |
| Lameness (chronic or recurrent) | 58 (5%) | 101 (11%) | 158 (11%) | 127 (11%) | 36 (7%) | 480 (9%) |
| Degenerative joint disease | 59 (5%) | 63 (7%) | 122 (8%) | 104 (9%) | 37 (7%) | 385 (7%) |
| Intervertebral disc disease (IVDD) | 167 (14%) | 77 (8%) | 41 (3%) | 20 (2%) | 13 (2%) | 318 (6%) |
| Elbow dysplasia | 23 (2%) | 20 (2%) | 50 (3%) | 61 (5%) | 30 (6%) | 184 (3%) |
| Rheumatoid arthritis | 35 (3%) | 34 (4%) | 47 (3%) | 47 (4%) | 17 (3%) | 180 (3%) |
| Spondylosis | 10 (<1%) | 29 (3%) | 46 (3%) | 28 (2%) | 9 (2%) | 122 (2%) |
| Growth deformity | 10 (<1%) | 7 (<1%) | 10 (<1%) | 7 (<1%) | 8 (2%) | 42 (<1%) |
| Panosteitis | 0 (<1%) | 2 (<1%) | 8 (<1%) | 20 (2%) | 11 (2%) | 41 (<1%) |
| Osteochondritis dissecans (OCD) | 0 (<1%) | 6 (<1%) | 8 (<1%) | 10 (<1%) | 14 (3%) | 38 (<1%) |
| Carpal subluxation syndrome | 7 (<1%) | 3 (<1%) | 5 (<1%) | 4 (<1%) | 0 (<1%) | 19 (<1%) |
| Dwarfism | 3 (<1%) | 5 (<1%) | 1 (<1%) | 0 (<1%) | 0 (<1%) | 9 (<1%) |
| Osteomyelitis | 1 (<1%) | 0 (<1%) | 1 (<1%) | 4 (<1%) | 2 (<1%) | 8 (<1%) |

Table 4: Proportion with Gastrointestinal Disorder History by Weight Category

|  | Weight Category | | | | |  |
| --- | --- | --- | --- | --- | --- | --- |
| Gastrointestinal | <10kg (N=892) | 10 to <20 (N=786) | 20 to <30 (N=1119) | 30 to <40 (N=749) | >=40 (N=368) | Total (N=3914) |
| Chronic or recurrent diarrhea | 150 (17%) | 168 (21%) | 217 (19%) | 163 (22%) | 88 (24%) | 786 (20%) |
| Anal sac impaction | 210 (24%) | 157 (20%) | 194 (17%) | 108 (14%) | 51 (14%) | 720 (18%) |
| Foreign body ingestion or blockage | 112 (13%) | 110 (14%) | 213 (19%) | 178 (24%) | 72 (20%) | 685 (18%) |
| Food or medicine allergies | 122 (14%) | 127 (16%) | 184 (16%) | 137 (18%) | 76 (21%) | 646 (17%) |
| Other | 102 (11%) | 92 (12%) | 113 (10%) | 81 (11%) | 31 (8%) | 419 (11%) |
| Irritable bowel syndrome (IBS or inflammatory bowel disease IBD) | 98 (11%) | 73 (9%) | 100 (9%) | 55 (7%) | 34 (9%) | 360 (9%) |
| Chronic or recurrent vomiting | 103 (12%) | 69 (9%) | 88 (8%) | 49 (7%) | 17 (5%) | 326 (8%) |
| Hemorrhagic gastroenteritis (HGE or stress colitis acute) | 71 (8%) | 56 (7%) | 56 (5%) | 45 (6%) | 20 (5%) | 248 (6%) |
| Other allergies | 28 (3%) | 20 (3%) | 36 (3%) | 17 (2%) | 14 (4%) | 115 (3%) |
| Bilious vomiting syndrome | 26 (3%) | 28 (4%) | 27 (2%) | 14 (2%) | 7 (2%) | 102 (3%) |
| Fecal incontinence | 12 (1%) | 21 (3%) | 31 (3%) | 16 (2%) | 7 (2%) | 87 (2%) |
| Constipation | 24 (3%) | 18 (2%) | 24 (2%) | 9 (1%) | 3 (<1%) | 78 (2%) |
| Bloat with torsion (GDV) | 1 (<1%) | 6 (<1%) | 28 (3%) | 20 (3%) | 10 (3%) | 65 (2%) |
| Idiopathic canine colitis (chronic) | 8 (<1%) | 11 (1%) | 15 (1%) | 6 (<1%) | 6 (2%) | 46 (1%) |
| Megaesophagus | 4 (<1%) | 0 (<1%) | 6 (<1%) | 6 (<1%) | 5 (1%) | 21 (<1%) |
| Protein losing enteropathy (PLE) | 7 (<1%) | 3 (<1%) | 3 (<1%) | 2 (<1%) | 0 (<1%) | 15 (<1%) |
| Malabsorptive disorder | 3 (<1%) | 2 (<1%) | 4 (<1%) | 1 (<1%) | 2 (<1%) | 12 (<1%) |
| Lymphangiectasia | 4 (<1%) | 2 (<1%) | 3 (<1%) | 0 (<1%) | 1 (<1%) | 10 (<1%) |
| Pyloric stenosis | 0 (<1%) | 0 (<1%) | 2 (<1%) | 0 (<1%) | 0 (<1%) | 2 (<1%) |

Table 5: Proportion with Eye Disorder History by Weight Category

|  | Weight Category | | | | |  |
| --- | --- | --- | --- | --- | --- | --- |
| Eye | <10kg (N=1079) | 10 to <20 (N=800) | 20 to <30 (N=934) | 30 to <40 (N=571) | >=40 (N=241) | Total (N=3625) |
| Adult onset cataracts | 470 (44%) | 238 (30%) | 245 (26%) | 123 (22%) | 38 (16%) | 1114 (31%) |
| Conjunctivitis | 117 (11%) | 173 (22%) | 275 (29%) | 138 (24%) | 56 (23%) | 759 (21%) |
| Other | 142 (13%) | 115 (14%) | 142 (15%) | 116 (20%) | 44 (18%) | 559 (15%) |
| Corneal ulcer | 119 (11%) | 75 (9%) | 66 (7%) | 50 (9%) | 11 (5%) | 321 (9%) |
| Dry eye (KCS) | 134 (12%) | 78 (10%) | 50 (5%) | 18 (3%) | 7 (3%) | 287 (8%) |
| Blindness (acquired) | 108 (10%) | 53 (7%) | 52 (6%) | 21 (4%) | 3 (1%) | 237 (7%) |
| Nuclear sclerosis (whitening of the eye) | 73 (7%) | 54 (7%) | 55 (6%) | 26 (5%) | 3 (1%) | 211 (6%) |
| Third eyelid prolapse (cherry eye) | 65 (6%) | 61 (8%) | 29 (3%) | 20 (4%) | 20 (8%) | 195 (5%) |
| Entropion (eyelid rolled in) | 7 (<1%) | 18 (2%) | 61 (7%) | 33 (6%) | 54 (22%) | 173 (5%) |
| Glaucoma | 47 (4%) | 29 (4%) | 24 (3%) | 12 (2%) | 5 (2%) | 117 (3%) |
| Uveitis | 13 (1%) | 12 (2%) | 25 (3%) | 17 (3%) | 3 (1%) | 70 (2%) |
| Progressive retinal atrophy or degeneration | 22 (2%) | 14 (2%) | 16 (2%) | 13 (2%) | 2 (<1%) | 67 (2%) |
| Juvenile cataracts | 13 (1%) | 9 (1%) | 15 (2%) | 16 (3%) | 8 (3%) | 61 (2%) |
| Iris cyst | 3 (<1%) | 7 (<1%) | 10 (1%) | 19 (3%) | 8 (3%) | 47 (1%) |
| Distichia | 12 (1%) | 7 (<1%) | 17 (2%) | 9 (2%) | 0 (<1%) | 45 (1%) |
| Ectropion (eyelid rolled out) | 8 (<1%) | 3 (<1%) | 10 (1%) | 9 (2%) | 14 (6%) | 44 (1%) |
| Retinal detachment | 15 (1%) | 10 (1%) | 6 (<1%) | 2 (<1%) | 1 (<1%) | 34 (<1%) |
| Pigmentary uveitis | 1 (<1%) | 0 (<1%) | 7 (<1%) | 9 (2%) | 2 (<1%) | 19 (<1%) |
| Imperforate lacrimal punctum | 4 (<1%) | 1 (<1%) | 2 (<1%) | 0 (<1%) | 0 (<1%) | 7 (<1%) |

Table 6: Proportion with Ear, Nose, and Throat Disorder History by Weight Category

|  | Weight Category | | | | |  |
| --- | --- | --- | --- | --- | --- | --- |
| Ear, Nose, and Throat | <10kg (N=827) | 10 to <20 (N=713) | 20 to <30 (N=915) | 30 to <40 (N=752) | >=40 (N=362) | Total (N=3569) |
| Chronic or recurrent ear infections | 312 (38%) | 334 (47%) | 534 (58%) | 538 (72%) | 280 (77%) | 1998 (56%) |
| Hearing loss (incompletely deaf) | 252 (30%) | 177 (25%) | 140 (15%) | 68 (9%) | 16 (4%) | 653 (18%) |
| Deafness (acquired) | 175 (21%) | 102 (14%) | 80 (9%) | 40 (5%) | 8 (2%) | 405 (11%) |
| Other | 72 (9%) | 64 (9%) | 76 (8%) | 69 (9%) | 27 (7%) | 308 (9%) |
| Ear mites | 71 (9%) | 62 (9%) | 77 (8%) | 48 (6%) | 31 (9%) | 289 (8%) |
| Hematoma | 5 (<1%) | 20 (3%) | 66 (7%) | 51 (7%) | 18 (5%) | 160 (4%) |
| Rhinitis | 27 (3%) | 17 (2%) | 26 (3%) | 9 (1%) | 3 (<1%) | 82 (2%) |
| Epistaxis (nose bleeds) | 1 (<1%) | 1 (<1%) | 10 (1%) | 4 (<1%) | 4 (1%) | 20 (<1%) |
| Tonsillitis | 4 (<1%) | 5 (<1%) | 5 (<1%) | 0 (<1%) | 2 (<1%) | 16 (<1%) |
| Pharyngitis | 3 (<1%) | 3 (<1%) | 6 (<1%) | 1 (<1%) | 1 (<1%) | 14 (<1%) |

Table 7: Proportion with Kidney or Urinary Disorder History by Weight Category

|  | Weight Category | | | | |  |
| --- | --- | --- | --- | --- | --- | --- |
| Kidney | <10kg (N=530) | 10 to <20 (N=452) | 20 to <30 (N=640) | 30 to <40 (N=369) | >=40 (N=131) | Total (N=2122) |
| Urinary tract infection (chronic or recurrent) | 173 (33%) | 161 (36%) | 267 (42%) | 155 (42%) | 57 (44%) | 813 (38%) |
| Urinary incontinence | 96 (18%) | 133 (29%) | 232 (36%) | 130 (35%) | 45 (34%) | 636 (30%) |
| Urinary crystals or stones in bladder or urethra | 176 (33%) | 95 (21%) | 87 (14%) | 53 (14%) | 12 (9%) | 423 (20%) |
| Chronic kidney disease | 80 (15%) | 53 (12%) | 52 (8%) | 24 (7%) | 5 (4%) | 214 (10%) |
| Other | 57 (11%) | 39 (9%) | 60 (9%) | 32 (9%) | 14 (11%) | 202 (10%) |
| Acute kidney failure | 24 (5%) | 8 (2%) | 20 (3%) | 8 (2%) | 8 (6%) | 68 (3%) |
| Proteinuria | 18 (3%) | 14 (3%) | 17 (3%) | 9 (2%) | 4 (3%) | 62 (3%) |
| Kidney stones | 22 (4%) | 8 (2%) | 9 (1%) | 2 (<1%) | 0 (<1%) | 41 (2%) |
| Pyelonephritis (kidney infection) | 3 (<1%) | 4 (<1%) | 4 (<1%) | 4 (1%) | 0 (<1%) | 15 (<1%) |
| Renal dysplasia | 2 (<1%) | 0 (<1%) | 3 (<1%) | 1 (<1%) | 0 (<1%) | 6 (<1%) |
| Ectopic ureter | 0 (<1%) | 2 (<1%) | 3 (<1%) | 0 (<1%) | 0 (<1%) | 5 (<1%) |
| Bladder prolapse | 1 (<1%) | 1 (<1%) | 1 (<1%) | 0 (<1%) | 0 (<1%) | 3 (<1%) |
| Urethral prolapse | 0 (<1%) | 0 (<1%) | 2 (<1%) | 0 (<1%) | 1 (<1%) | 3 (<1%) |
| Tubular disorder (such as Fanconi syndrome) | 2 (<1%) | 0 (<1%) | 0 (<1%) | 0 (<1%) | 0 (<1%) | 2 (<1%) |

Table 8-1: Proportion with Cancer or Tumors History by Weight Category (Areas of Body)

|  | Weight Category | | | | |  |
| --- | --- | --- | --- | --- | --- | --- |
| Cancer (Areas of Body) | <10kg (N=273) | 10 to <20 (N=327) | 20 to <30 (N=565) | 30 to <40 (N=428) | >=40 (N=158) | Total (N=1751) |
| Skin of trunk, body, or head | 61 (22%) | 75 (23%) | 186 (33%) | 136 (32%) | 44 (28%) | 502 (29%) |
| Muscle or other soft tissue | 55 (20%) | 61 (19%) | 118 (21%) | 106 (25%) | 36 (23%) | 376 (21%) |
| Skin of limb or foot | 30 (11%) | 39 (12%) | 99 (18%) | 61 (14%) | 25 (16%) | 254 (15%) |
| Other location of cancer | 15 (5%) | 33 (10%) | 37 (7%) | 36 (8%) | 7 (4%) | 128 (7%) |
| Oral (mouth) cavity | 15 (5%) | 23 (7%) | 32 (6%) | 29 (7%) | 9 (6%) | 108 (6%) |
| Spleen | 7 (3%) | 24 (7%) | 35 (6%) | 18 (4%) | 7 (4%) | 91 (5%) |
| Bone or joint | 8 (3%) | 8 (2%) | 20 (4%) | 26 (6%) | 21 (13%) | 83 (5%) |
| Mammary (breast) tissue | 27 (10%) | 12 (4%) | 20 (4%) | 9 (2%) | 6 (4%) | 74 (4%) |
| Lymph nodes | 7 (3%) | 17 (5%) | 13 (2%) | 28 (7%) | 6 (4%) | 71 (4%) |
| Liver | 10 (4%) | 11 (3%) | 20 (4%) | 11 (3%) | 1 (<1%) | 53 (3%) |
| Anal sac | 9 (3%) | 12 (4%) | 14 (2%) | 13 (3%) | 1 (<1%) | 49 (3%) |
| Eye | 12 (4%) | 8 (2%) | 12 (2%) | 10 (2%) | 4 (3%) | 46 (3%) |
| Ear | 9 (3%) | 7 (2%) | 15 (3%) | 10 (2%) | 4 (3%) | 45 (3%) |
| Lung | 5 (2%) | 8 (2%) | 7 (1%) | 12 (3%) | 2 (1%) | 34 (2%) |
| Perianal area | 7 (3%) | 6 (2%) | 10 (2%) | 6 (1%) | 0 (<1%) | 29 (2%) |
| Dont know | 3 (1%) | 5 (2%) | 11 (2%) | 7 (2%) | 2 (1%) | 28 (2%) |
| Gastrointestinal tract (stomach and/or intestine) | 4 (1%) | 7 (2%) | 8 (1%) | 4 (<1%) | 3 (2%) | 26 (1%) |
| Adrenal gland | 7 (3%) | 6 (2%) | 7 (1%) | 2 (<1%) | 1 (<1%) | 23 (1%) |
| Bladder or urethra | 8 (3%) | 9 (3%) | 6 (1%) | 0 (<1%) | 0 (<1%) | 23 (1%) |
| Rectum | 2 (<1%) | 4 (1%) | 10 (2%) | 4 (<1%) | 2 (1%) | 22 (1%) |
| Thyroid | 7 (3%) | 1 (<1%) | 7 (1%) | 4 (<1%) | 0 (<1%) | 19 (1%) |
| Nose or nasal passage | 3 (1%) | 3 (<1%) | 7 (1%) | 4 (<1%) | 1 (<1%) | 18 (1%) |
| Nerve sheath | 1 (<1%) | 5 (2%) | 4 (<1%) | 5 (1%) | 3 (2%) | 18 (1%) |
| Blood | 3 (1%) | 2 (<1%) | 5 (<1%) | 6 (1%) | 1 (<1%) | 17 (<1%) |
| Brain | 1 (<1%) | 3 (<1%) | 4 (<1%) | 3 (<1%) | 1 (<1%) | 12 (<1%) |
| Kidney | 4 (1%) | 0 (<1%) | 2 (<1%) | 4 (<1%) | 2 (1%) | 12 (<1%) |
| Testicle | 1 (<1%) | 2 (<1%) | 0 (<1%) | 3 (<1%) | 3 (2%) | 9 (<1%) |
| Venereal (vagina, labia, penis, prepuce) | 2 (<1%) | 2 (<1%) | 2 (<1%) | 3 (<1%) | 0 (<1%) | 9 (<1%) |
| Pancreas | 3 (1%) | 0 (<1%) | 5 (<1%) | 0 (<1%) | 0 (<1%) | 8 (<1%) |
| Pituitary gland | 0 (<1%) | 1 (<1%) | 6 (1%) | 0 (<1%) | 0 (<1%) | 7 (<1%) |
| Spinal cord | 1 (<1%) | 2 (<1%) | 3 (<1%) | 0 (<1%) | 1 (<1%) | 7 (<1%) |
| Cardiac (heart) tissue | 0 (<1%) | 1 (<1%) | 3 (<1%) | 2 (<1%) | 0 (<1%) | 6 (<1%) |
| Gallbladder or bile duct | 0 (<1%) | 1 (<1%) | 2 (<1%) | 2 (<1%) | 0 (<1%) | 5 (<1%) |
| Ovary or uterus | 1 (<1%) | 3 (<1%) | 1 (<1%) | 0 (<1%) | 0 (<1%) | 5 (<1%) |
| Prostate | 0 (<1%) | 3 (<1%) | 1 (<1%) | 0 (<1%) | 0 (<1%) | 4 (<1%) |
| Esophagus | 0 (<1%) | 0 (<1%) | 1 (<1%) | 0 (<1%) | 0 (<1%) | 1 (<1%) |

Table 8-2: Proportion with Cancer or Tumors History by Weight Category (Types of Cancer)

|  | Weight Category | | | | |  |
| --- | --- | --- | --- | --- | --- | --- |
| Cancer (Types of Cancer) | <10kg (N=273) | 10 to <20 (N=327) | 20 to <30 (N=565) | 30 to <40 (N=428) | >=40 (N=158) | Total (N=1751) |
| Dont know | 95 (35%) | 112 (34%) | 124 (22%) | 117 (27%) | 51 (32%) | 499 (28%) |
| Mast cell tumor | 39 (14%) | 50 (15%) | 140 (25%) | 97 (23%) | 23 (15%) | 349 (20%) |
| Lipoma | 29 (11%) | 40 (12%) | 76 (13%) | 52 (12%) | 20 (13%) | 217 (12%) |
| Other type of cancer | 23 (8%) | 30 (9%) | 42 (7%) | 30 (7%) | 16 (10%) | 141 (8%) |
| Soft tissue sarcoma | 7 (3%) | 14 (4%) | 29 (5%) | 17 (4%) | 7 (4%) | 74 (4%) |
| Melanoma | 9 (3%) | 7 (2%) | 24 (4%) | 15 (4%) | 7 (4%) | 62 (4%) |
| Carcinoma (not listed elsewhere) | 9 (3%) | 10 (3%) | 17 (3%) | 15 (4%) | 6 (4%) | 57 (3%) |
| Lymphoma lymphosarcoma | 6 (2%) | 8 (2%) | 7 (1%) | 25 (6%) | 4 (3%) | 50 (3%) |
| Sarcoma (not listed elsewhere) | 4 (1%) | 7 (2%) | 19 (3%) | 15 (4%) | 2 (1%) | 47 (3%) |
| Adenoma (not listed elsewhere) | 10 (4%) | 10 (3%) | 15 (3%) | 7 (2%) | 1 (<1%) | 43 (2%) |
| Adenocarcinoma (not listed elsewhere) | 6 (2%) | 11 (3%) | 17 (3%) | 8 (2%) | 0 (<1%) | 42 (2%) |
| Hemangiosarcoma | 2 (<1%) | 7 (2%) | 18 (3%) | 12 (3%) | 2 (1%) | 41 (2%) |
| Histiocytoma | 5 (2%) | 8 (2%) | 17 (3%) | 5 (1%) | 3 (2%) | 38 (2%) |
| Basal cell tumor | 8 (3%) | 6 (2%) | 10 (2%) | 9 (2%) | 2 (1%) | 35 (2%) |
| Osteosarcoma | 3 (1%) | 4 (1%) | 8 (1%) | 6 (1%) | 9 (6%) | 30 (2%) |
| Epidermoid cyst | 4 (1%) | 4 (1%) | 11 (2%) | 7 (2%) | 3 (2%) | 29 (2%) |
| Hemangioma | 2 (<1%) | 5 (2%) | 10 (2%) | 5 (1%) | 2 (1%) | 24 (1%) |
| Squamous cell carcinoma | 3 (1%) | 4 (1%) | 7 (1%) | 5 (1%) | 2 (1%) | 21 (1%) |
| Papilloma | 2 (<1%) | 4 (1%) | 12 (2%) | 2 (<1%) | 0 (<1%) | 20 (1%) |
| Sebaceous adenoma | 3 (1%) | 3 (<1%) | 3 (<1%) | 7 (2%) | 0 (<1%) | 16 (<1%) |
| Fibrosarcoma | 4 (1%) | 1 (<1%) | 8 (1%) | 1 (<1%) | 1 (<1%) | 15 (<1%) |
| Plasmacytoma | 1 (<1%) | 2 (<1%) | 6 (1%) | 5 (1%) | 1 (<1%) | 15 (<1%) |
| Epulides | 0 (<1%) | 3 (<1%) | 6 (1%) | 4 (<1%) | 1 (<1%) | 14 (<1%) |
| Peripheral nerve sheath tumor | 1 (<1%) | 3 (<1%) | 3 (<1%) | 3 (<1%) | 2 (1%) | 12 (<1%) |
| Transitional cell carcinoma | 3 (1%) | 4 (1%) | 3 (<1%) | 0 (<1%) | 0 (<1%) | 10 (<1%) |
| Leukemia | 2 (<1%) | 0 (<1%) | 1 (<1%) | 3 (<1%) | 0 (<1%) | 6 (<1%) |
| Histiocytic sarcoma | 0 (<1%) | 0 (<1%) | 1 (<1%) | 2 (<1%) | 2 (1%) | 5 (<1%) |
| Insulinoma | 1 (<1%) | 0 (<1%) | 3 (<1%) | 1 (<1%) | 0 (<1%) | 5 (<1%) |
| Chondrosarcoma | 1 (<1%) | 1 (<1%) | 1 (<1%) | 1 (<1%) | 0 (<1%) | 4 (<1%) |
| Cystadenoma | 1 (<1%) | 0 (<1%) | 1 (<1%) | 0 (<1%) | 0 (<1%) | 2 (<1%) |
| Leiomyoma | 1 (<1%) | 0 (<1%) | 0 (<1%) | 0 (<1%) | 1 (<1%) | 2 (<1%) |
| Meningioma | 1 (<1%) | 0 (<1%) | 1 (<1%) | 0 (<1%) | 0 (<1%) | 2 (<1%) |
| Leiomyosarcoma | 0 (<1%) | 0 (<1%) | 1 (<1%) | 0 (<1%) | 0 (<1%) | 1 (<1%) |
| Multiple myeloma | 1 (<1%) | 0 (<1%) | 0 (<1%) | 0 (<1%) | 0 (<1%) | 1 (<1%) |
| Thymoma | 0 (<1%) | 0 (<1%) | 0 (<1%) | 1 (<1%) | 0 (<1%) | 1 (<1%) |
| Rhabdomyosarcoma | 0 (<1%) | 0 (<1%) | 0 (<1%) | 0 (<1%) | 0 (<1%) | 0 (<1%) |

Table 9: Proportion with Cardiac Disorder History by Weight Category

|  | Weight Category | | | | |  |
| --- | --- | --- | --- | --- | --- | --- |
| Cardiac | <10kg (N=671) | 10 to <20 (N=362) | 20 to <30 (N=301) | 30 to <40 (N=177) | >=40 (N=56) | Total (N=1567) |
| Murmur | 541 (81%) | 269 (74%) | 195 (65%) | 116 (66%) | 35 (62%) | 1156 (74%) |
| Valve disease | 63 (9%) | 33 (9%) | 25 (8%) | 14 (8%) | 4 (7%) | 139 (9%) |
| Congestive heart failure | 84 (13%) | 29 (8%) | 12 (4%) | 10 (6%) | 2 (4%) | 137 (9%) |
| Arrhythmia | 33 (5%) | 24 (7%) | 32 (11%) | 23 (13%) | 12 (21%) | 124 (8%) |
| Cardiomyopathy | 19 (3%) | 17 (5%) | 30 (10%) | 22 (12%) | 5 (9%) | 93 (6%) |
| Other | 28 (4%) | 18 (5%) | 22 (7%) | 11 (6%) | 1 (2%) | 80 (5%) |
| Hypertension (high blood pressure) | 29 (4%) | 26 (7%) | 13 (4%) | 10 (6%) | 0 (<1%) | 78 (5%) |
| Pulmonary hypertension | 15 (2%) | 2 (<1%) | 4 (1%) | 2 (1%) | 0 (<1%) | 23 (1%) |
| Subaortic stenosis | 0 (<1%) | 1 (<1%) | 9 (3%) | 6 (3%) | 3 (5%) | 19 (1%) |
| Pulmonic stenosis | 2 (<1%) | 3 (<1%) | 2 (<1%) | 1 (<1%) | 2 (4%) | 10 (<1%) |
| Endocarditis | 3 (<1%) | 1 (<1%) | 2 (<1%) | 1 (<1%) | 0 (<1%) | 7 (<1%) |
| Pericardial effusion | 2 (<1%) | 0 (<1%) | 0 (<1%) | 0 (<1%) | 0 (<1%) | 2 (<1%) |

Table 10: Proportion with Neurologic Disorder History by Weight Category

|  | Weight Category | | | | |  |
| --- | --- | --- | --- | --- | --- | --- |
| Neurologic | <10kg (N=362) | 10 to <20 (N=278) | 20 to <30 (N=346) | 30 to <40 (N=235) | >=40 (N=103) | Total (N=1324) |
| Seizures (including epilepsy) | 186 (51%) | 135 (49%) | 139 (40%) | 100 (43%) | 46 (45%) | 606 (46%) |
| Other | 55 (15%) | 47 (17%) | 66 (19%) | 45 (19%) | 18 (17%) | 231 (17%) |
| Dementia or senility | 59 (16%) | 39 (14%) | 30 (9%) | 18 (8%) | 7 (7%) | 153 (12%) |
| Vestibular disease | 23 (6%) | 35 (13%) | 63 (18%) | 25 (11%) | 1 (<1%) | 147 (11%) |
| Intervertebral disc disease (IVDD 1 | 52 (14%) | 23 (8%) | 20 (6%) | 4 (2%) | 3 (3%) | 102 (8%) |
| Laryngeal paralysis | 1 (<1%) | 3 (1%) | 28 (8%) | 20 (9%) | 13 (13%) | 65 (5%) |
| Degenerative myelopathy | 7 (2%) | 12 (4%) | 15 (4%) | 17 (7%) | 8 (8%) | 59 (4%) |
| Limb paralysis | 14 (4%) | 5 (2%) | 8 (2%) | 5 (2%) | 4 (4%) | 36 (3%) |
| Horners syndrome | 1 (<1%) | 3 (1%) | 9 (3%) | 12 (5%) | 2 (2%) | 27 (2%) |
| Wobbler syndrome | 4 (1%) | 5 (2%) | 1 (<1%) | 3 (1%) | 5 (5%) | 18 (1%) |
| Polyneuropathy | 2 (<1%) | 2 (<1%) | 4 (1%) | 3 (1%) | 4 (4%) | 15 (1%) |
| Fibrocartilaginous embolism (FCE) | 1 (<1%) | 3 (1%) | 2 (<1%) | 1 (<1%) | 4 (4%) | 11 (<1%) |
| Cauda equina syndrome | 1 (<1%) | 1 (<1%) | 2 (<1%) | 3 (1%) | 1 (<1%) | 8 (<1%) |
| Diskospondylitis | 1 (<1%) | 1 (<1%) | 0 (<1%) | 0 (<1%) | 2 (2%) | 4 (<1%) |
| Myasthenia gravis | 0 (<1%) | 0 (<1%) | 1 (<1%) | 2 (<1%) | 1 (<1%) | 4 (<1%) |
| Dysautonomia | 0 (<1%) | 0 (<1%) | 0 (<1%) | 0 (<1%) | 0 (<1%) | 0 (<1%) |

Table 11: Proportion with Liver or Pancreas Disorder History by Weight Category

|  | Weight Category | | | | |  |
| --- | --- | --- | --- | --- | --- | --- |
| Liver | <10kg (N=393) | 10 to <20 (N=209) | 20 to <30 (N=195) | 30 to <40 (N=130) | >=40 (N=43) | Total (N=970) |
| Pancreatitis | 245 (62%) | 126 (60%) | 110 (56%) | 77 (59%) | 29 (67%) | 587 (61%) |
| Other | 82 (21%) | 49 (23%) | 50 (26%) | 26 (20%) | 9 (21%) | 216 (22%) |
| Chronic inflammatory liver disease | 37 (9%) | 20 (10%) | 20 (10%) | 9 (7%) | 0 (<1%) | 86 (9%) |
| Exocrine pancreatic insufficiency (EPI) | 8 (2%) | 5 (2%) | 7 (4%) | 11 (8%) | 4 (9%) | 35 (4%) |
| Gall bladder mucocele | 19 (5%) | 12 (6%) | 2 (1%) | 1 (<1%) | 1 (2%) | 35 (4%) |
| Gall bladder surgery | 5 (1%) | 4 (2%) | 2 (1%) | 1 (<1%) | 1 (2%) | 13 (1%) |
| Microvascular dysplasia (portal vein hypoplasia) | 8 (2%) | 0 (<1%) | 0 (<1%) | 0 (<1%) | 0 (<1%) | 8 (<1%) |
| Biliary obstruction | 2 (<1%) | 2 (<1%) | 2 (1%) | 0 (<1%) | 1 (2%) | 7 (<1%) |
| Gall bladder rupture | 3 (<1%) | 1 (<1%) | 2 (1%) | 0 (<1%) | 0 (<1%) | 6 (<1%) |
| Portosystemic shunt (acquired) | 1 (<1%) | 1 (<1%) | 1 (<1%) | 1 (<1%) | 0 (<1%) | 4 (<1%) |

Table 12: Proportion with Respiratory Disorder History by Weight Category

|  | Weight Category | | | | |  |
| --- | --- | --- | --- | --- | --- | --- |
| Respiratory | <10kg (N=385) | 10 to <20 (N=169) | 20 to <30 (N=207) | 30 to <40 (N=124) | >=40 (N=65) | Total (N=950) |
| Chronic or recurrent cough | 122 (32%) | 48 (28%) | 39 (19%) | 19 (15%) | 8 (12%) | 236 (25%) |
| Tracheal collapse | 147 (38%) | 15 (9%) | 4 (2%) | 2 (2%) | 1 (2%) | 169 (18%) |
| Pneumonia | 19 (5%) | 42 (25%) | 51 (25%) | 28 (23%) | 23 (35%) | 163 (17%) |
| Laryngeal paralysis | 3 (<1%) | 6 (4%) | 44 (21%) | 37 (30%) | 23 (35%) | 113 (12%) |
| Other | 31 (8%) | 20 (12%) | 38 (18%) | 14 (11%) | 5 (8%) | 108 (11%) |
| Tracheal stenosis (narrowing ) | 57 (15%) | 7 (4%) | 6 (3%) | 7 (6%) | 3 (5%) | 80 (8%) |
| Chronic or recurrent bronchitis | 26 (7%) | 17 (10%) | 13 (6%) | 9 (7%) | 3 (5%) | 68 (7%) |
| Elongated soft palate | 20 (5%) | 17 (10%) | 15 (7%) | 3 (2%) | 0 (<1%) | 55 (6%) |
| Chronic or recurrent rhinitis | 14 (4%) | 9 (5%) | 14 (7%) | 5 (4%) | 5 (8%) | 47 (5%) |
| Stenotic narrow nares | 19 (5%) | 15 (9%) | 6 (3%) | 0 (<1%) | 0 (<1%) | 40 (4%) |
| Acquired or acute respiratory distress syndrome (ARDS) | 1 (<1%) | 0 (<1%) | 5 (2%) | 2 (2%) | 1 (2%) | 9 (<1%) |
| Pulmonary bullae | 0 (<1%) | 0 (<1%) | 2 (<1%) | 2 (2%) | 1 (2%) | 5 (<1%) |
| Lung lobe torsion | 0 (<1%) | 0 (<1%) | 1 (<1%) | 0 (<1%) | 0 (<1%) | 1 (<1%) |

Table 13: Proportion with Endocrine Disorder History by Weight Category

|  | Weight Category | | | | |  |
| --- | --- | --- | --- | --- | --- | --- |
| Endocrine | <10kg (N=194) | 10 to <20 (N=190) | 20 to <30 (N=264) | 30 to <40 (N=176) | >=40 (N=89) | Total (N=913) |
| Hypothyroidism (low thyroid function) | 78 (40%) | 86 (45%) | 172 (65%) | 118 (67%) | 66 (74%) | 520 (57%) |
| Cushings disease (hyperadrenocorticism excess adrenal function) | 65 (34%) | 55 (29%) | 35 (13%) | 15 (9%) | 5 (6%) | 175 (19%) |
| Diabetes mellitus (common diabetes which causes high blood sugar) | 37 (19%) | 22 (12%) | 11 (4%) | 11 (6%) | 2 (2%) | 83 (9%) |
| Addisons disease (hypoadrenocorticism low adrenal function) | 14 (7%) | 15 (8%) | 25 (9%) | 14 (8%) | 8 (9%) | 76 (8%) |
| Diabetes insipidus (rare diabetes which causes water balance problems) | 4 (2%) | 5 (3%) | 14 (5%) | 6 (3%) | 2 (2%) | 31 (3%) |
| Hypoparathyroidism (low parathyroid function causing low calcium) | 4 (2%) | 6 (3%) | 3 (1%) | 6 (3%) | 5 (6%) | 24 (3%) |
| Hyperthyroidism (excess thyroid function) | 5 (3%) | 3 (2%) | 8 (3%) | 5 (3%) | 3 (3%) | 24 (3%) |
| Hyperparathyroidism (excess parathyroid function causing high calcium) | 2 (1%) | 4 (2%) | 5 (2%) | 2 (1%) | 0 (<1%) | 13 (1%) |
| Other | 3 (2%) | 4 (2%) | 2 (<1%) | 1 (<1%) | 0 (<1%) | 10 (1%) |
| Hypercalcemia (excess calcium in the blood) | 1 (<1%) | 1 (<1%) | 3 (1%) | 2 (1%) | 0 (<1%) | 7 (<1%) |
